# Universal MS/MS Visualization and Retrieval with the Metabolomics Spectrum Resolver Web Service

**DOI:** 10.1101/2020.05.09.086066

**Authors:** Wout Bittremieux, Christopher Chen, Pieter C. Dorrestein, Emma L. Schymanski, Tobias Schulze, Steffen Neumann, Rene Meier, Simon Rogers, Mingxun Wang

## Abstract

The growth of online mass spectrometry metabolomics resources, including data repositories, spectral library databases, and online analysis platforms has created an environment of online/web accessibility. Here, we introduce the Metabolomics Spectrum Resolver (https://metabolomics-usi.ucsd.edu/), a tool that builds upon these exciting developments to allow for consistent data export (in human and machine-readable forms) and publication-ready visualisations of tandem mass spectrometry spectra. This tool supports the Human Proteome Organization – Proteomics Standards Initiative’s Universal Spectrum Identifier (USI) specification, which has been extended to deal with the metabolomics use cases. To date, this resource already supports data formats from GNPS, MassBank, MS2LDA, MassIVE, MetaboLights, and Metabolomics Workbench and is integrated into several of these resources.

## Introduction

The effective exchange and visualization of tandem mass spectrometry (MS/MS) information across a variety of resources is important in communicating and consequently building confidence in both data quality and molecule identification throughout the research and publication process. The inclusion of MS/MS spectra in scientific manuscripts and presentations to provide evidence of newly discovered molecules in a consistent manner is often challenging and labor intensive due to heterogeneous data availability and accessibility across public data resources, the variety of complex file formats in use to store mass spectrometry data, and the lack of suitable software tools available. Furthermore, it is often difficult to balance the ease of MS/MS figure generation with the level of customization possible.

A few approaches have been introduced to assign universal identifiers to MS/MS spectra. The SPectraL hASH (SPLASH) identifier has been designed as an unambiguous, database-independent spectrum identifier^1^. SPLASH identifiers can be used to determine spectral overlap between libraries and potentially query public data repositories. However, as generating a SPLASH identifier includes a complex hashing operation, these identifiers cannot be trivially produced. Alternatively, the Universal Spectrum Identifier (USI) was recently developed by the Human Proteome Organization – Proteomics Standards Initiative (HUPO-PSI)^2^ to provide a standardized mechanism for encoding a virtual path to spectra contained in proteomics public repositories^3^. Although USIs are easy to construct, the USI standard currently mainly supports proteomics spectral data deposited via ProteomeXchange^4^ and does not natively support spectral data stored in metabolomics repositories.

Additionally, producing high-quality figures of MS/MS spectra is challenging as well. On one hand, existing software such as the Proteomics Data Viewer (PDV)^5^, the Interactive Peptide Spectral Annotator (IPSA)^6^, and Lorikeet^7^ proteomics data viewers, as well as vendor software, are able to draw MS/MS spectra, but are mainly aimed at interactive visualization. Three key features are often missing from such software: the ability to export vector graphics (to ensure high resolution), customization of spectral visualization, and high-throughput automated figure generation. On the other end of the spectrum are generic vector graphics editors such as Adobe Illustrator or Inkscape. While such generic editors are powerful, they lack MS/MS specific features and producing images ready for publication with these tools requires a significant time investment to achieve high levels of customization.

We present here the Metabolomics Spectrum Resolver (https://metabolomics-usi.ucsd.edu/), a web tool that enables: 1) integration with major metabolomics data repositories to retrieve MS/MS metabolomics data from various sources in a unified fashion, 2) high-quality and customizable vector graphics drawing, 3) direct integration with common online analysis tools, and 4) a web API to programmatically retrieve spectral data.

## Results and Discussion

### MS/MS Data Sources - Integration with Community Resources

The Metabolomics Spectrum Resolver builds upon the USI standard developed by the HUPO-PSI^3^. USIs are formatted as follows:

~~~
mzspec:<collection>:<msRun>:<indexType>:<indexNumber>:<optional interpretation>
~~~

For more details on the USI, including its formal specification, see the HUPO-PSI website (http://www.psidev.info/usi).

The USI standard has originally been developed for proteomics data, with only ProteomeXchange identifiers and related identifiers from its member repositories allowed in the “collection” field. We have extended the USI specification to support several major metabolomics resources as well. Concretely, the following metabolomics data services are currently supported (**Table 1**):

- Data repositories: MassIVE^8^, MetaboLights^9^, Metabolomics Workbench^10^.
- Reference spectral library resources: MassBank^11^, MoNA (https://mona.fiehnlab.ucdavis.edu/), GNPS^12^, and MS2LDA.org MOTIFDB^13^.
- Online informatics pipelines: GNPS Molecular Networking^12^ / Library Search / MASST^14^ / Feature-based Molecular Networking^15^, and MS2LDA.org^13^.

**Table 1 -.** Examples of the Metabolomics Spectrum Resolver from various public data resources.

| Resource | USI Format | Example USI |
| --- | --- | --- |
| MassBank | collection: MASSBANK<br>msRun: none<br>index: accession:<spectrumId> | mzspec:MASSBANK::accession:SM858102 |
| GNPS Spectral Library | collection: GNPS<br>msRun: GNPS-LIBRARY<br>index: accession:<spectrumId> | mzspec:GNPS:GNPS-LIBRARY:accession:CCMSLIB00005436077 |
| GNPS Molecular Networking Spectrum | collection: GNPS<br>msRun: TASK-<taskId><br>index: scan:<scanNr> | mzspec:GNPS:TASK-c95481f0c53d42e78a61bf899e9f9adb-spectra/specs_ms.mgf:scan:1943 |
| MS2LDA MotifDB | collection: MOTIFDB<br>msRun: none<br>index: accession:<motifId> | mzspec:MOTIFDB::accession:171163 |
| MS2LDA Analysis Spectrum | collection: MS2LDA<br>msRun: TASK-<taskId><br>Index: accession:<spectrumId> | mzspec:MS2LDA:TASK-190:accession:270684 |
| MassIVE/GNPS Repository Spectrum | collection:MassIVE/ProteomeXchange identifier<br>msRun: filename<br>index: scan:<scanNr> | mzspec:MSV000078547:120228_nbut_3610_it_it_take2:scan:389 |
| Metabolights Repository Spectrum | collection: MSV000082791<br>msRun: filename<br>index: scan:<scanNr> | mzspec:MSV000082791:(-)-epigallocatechin:scan:2 |
| Metabolomics Workbench Repository Spectrum | collection: MSV000082680<br>msRun: filename<br>index: scan:<scanNr> | mzspec:MSV000082680:iPSC-T1R1:scan:3 |

As such, the Metabolomics Spectrum Resolver provides a universal interface to more than 450M MS/MS spectra from various public metabolomics repositories. It also supports linking back to the source spectra in their original data resources to allow users to explore the original context of the MS/MS spectra. Several resources (MassBank, GNPS, and MS2LDA.org) have already integrated links to the Metabolomics Spectrum Resolver to facilitate bidirectional interoperability. Additionally, SPLASH identifiers^1^ are computed for all spectra retrieved through the Metabolomics Spectrum Resolver to enable their comparison across different resources.

### Visualization Capabilities

The Metabolomics Spectrum Resolver enables users to plot individual MS/MS spectra (**Figure 1a**) and mirror matches between pairs of MS/MS spectra to show their similarity (e.g. between an experimental spectrum and its spectral library match; **Figure 1b**).The resulting figures can be modified by specifying mass ranges, intensity ranges, customizable peak labeling, figure size, decimal points for mass values, and an optional grid. Additionally, the similarity between the two spectra in a mirror plot can be displayed based on the standard cosine similarity or a “shifted cosine similarity” (i.e. also matching peaks that differ by the precursor mass difference between both spectra), with matching peaks between the two spectra highlighted to assess the match quality. The resulting drawing can be downloaded in the SVG (vector), PNG (raster), CSV (comma separated peaks), and JSON (machine readable peaks) formats (**Figure 2**).

**Figure 1 -.**
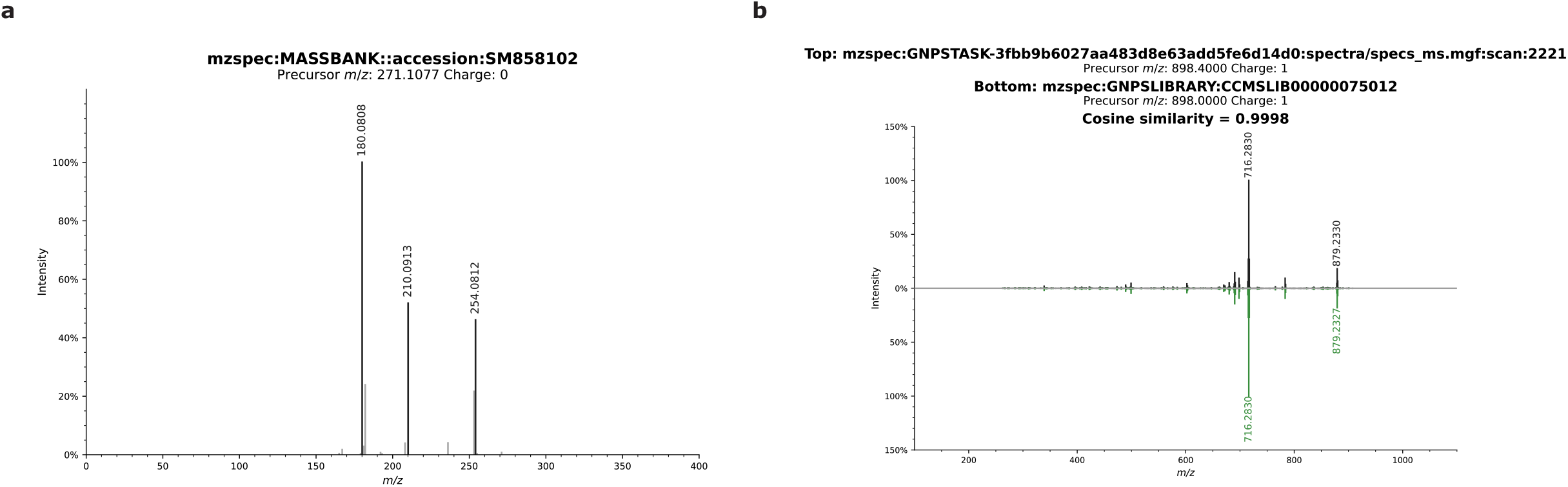
Example MS/MS. figure generation of (a) a single MS/MS <u>spectrum</u> and (b) a <u>mirror plot</u>.

**Figure 2 -.**
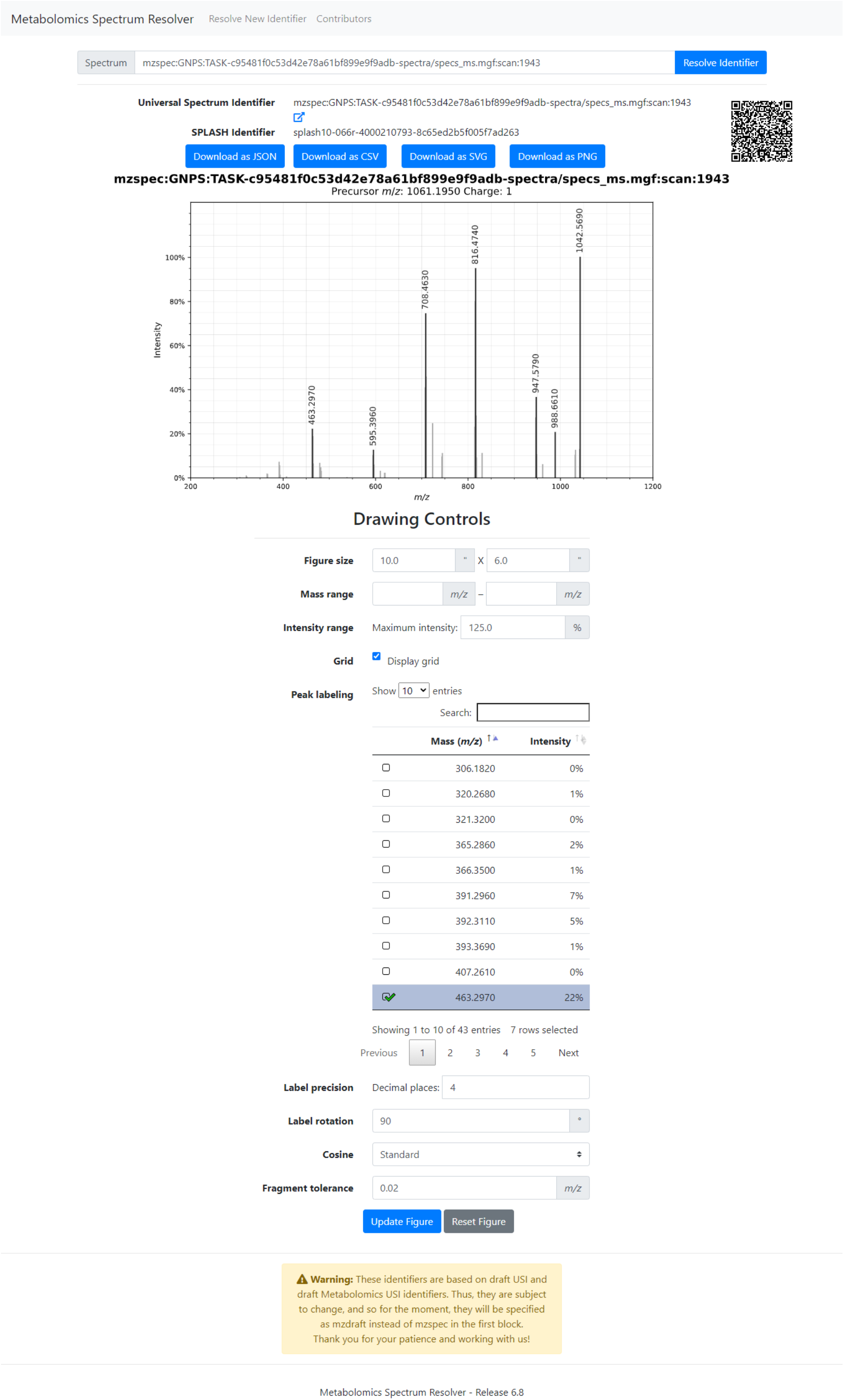
Interactive User Interface with Drawing Options. Enables customizing the spectrum drawing, linking back to original spectral data via URL and a generated QR code, the spectrum’s SPLASH identifier, spectrum peaks download, and programmatic data download.

Plotting functionality is implemented in Python using spectrum_utils^17^, Matplotlib^18^, and Seaborn^19^. Additionally, NumPy^20^, SciPy^21^, and Numba^22^ are used for efficient processing of spectral data.

## Conclusion

The Metabolomics Spectrum Resolver has been designed to improve the accessibility and presentation of metabolomics data, both within supported public data resources and for the community. By extending the nascent USI standard developed by the HUPO-PSI to support metabolomics repositories and tools as well, it provides a universal interface to metabolomics spectral data from heterogeneous data resources. The Metabolomics Spectral Resolver supports a broad range of key online metabolomics resources, provides unified programmatic API access to spectral data, and facilitates high quality spectrum drawing. Implemented as a freely available and open source web service, it facilitates and democratizes public data access without the need to install specialized software.

## Source Code

The Metabolomics Spectrum Resolver source code is released as open source under the MIT License and is available at this DOI: 10.5281/zenodo.4033442. Active development can be found on GitHub: https://github.com/mwang87/MetabolomicsSpectrumResolver.

## Acknowledgements

We thank Madeleine Ernst, Alan Jarmusch, and Louis-Felix Nothias for their feedback on the usability of the Metabolomics Spectrum Resolver. We thank Eric Deutsch and Nuno Bandeira for feedback on the metabolomics spectrum resolvers. We thank all past and future contributors to the resolver software. WB is a postdoctoral researcher of the Research Foundation – Flanders (FWO). ELS acknowledges funding support from the Luxembourg National Research Fund (FNR) for project A18/BM/12341006. MassBank Europe (https://massbank.eu/MassBank) is supported by the NORMAN Association (https://www.norman-network.net), de.NBI (FKZ 031L0107), NaToxAq (Marie Sklodowska-Curie grant agreement No. 722493), HBM4EU (European Union H2020 grant agreement No. 733032) and the Helmholtz Centre for Environmental Research - UFZ (https://www.ufz.de/index.php?en=33573).

